# Resting state EEG microstates: is a new FORM all you need?

**DOI:** 10.64898/2026.09.24.754021

**Authors:** Leonardo Corsi, Piergiuseppe Liuzzi, Calogero Maria Oddo, Andrea Mannini

## Abstract

**Introduction:** Before asking which microstate an EEG sample belongs to, there is a simpler question: can it look like a microstate at all? Yet a winning correlation mixes three different properties: how microstate-like the scalp field is, where it lies within that geometry, and how well the available templates represent that direction. We asked whether a fixed geometry, independent of any microstate solution, could explain common structure of microstate maps, constrain how strongly EEG patterns can match them, and describe that structure in interpretable terms.

**Methods:** We introduce First-Order spherical haRMonic (FORM), projecting normalized scalp topographies onto a fixed three-dimensional subspace defined only by sensor geometry. Direction describes field geometry; projection norm, ρ, measures FORM conformity and the maximum correlation attainable by any FORM topography. We tested simulations, literature templates, and 374 recordings from 187 LEMON participants.

**Results:** FORM captured canonical microstate geometry despite being defined a priori of a microstate solution: FORM explained energy ρ^2^ was 0.990-0.998 for five meta-microstates and 0.966-0.986 for pooled LEMON maps. Across 313 literature templates, FORM angular distances preserved their independently derived organization (Spearman *ρ* = 0.987). In continuous EEG, FORM conformity explained a median 74.4% of within-subject variance in winning-template correlation, versus 21.7% for GFP. As independently estimated dictionaries increased from *K* = 1 to *K* = 12, the median fitted fraction of the FORM ceiling rose from 0.613 to 0.970.

**Discussion:** FORM places an a priori continuous geometry beneath discrete microstate labels. It separates properties intrinsic to the scalp field - whether it is microstate-like and where it lies within that geometry - from how well the chosen template dictionary represents it. FORM therefore provides an interpretable, clustering-independent reference for assignment strength, template comparison and continuous microstate dynamics.

## 1. Introduction

In microstate analysis, resting-state EEG is described as a sequence of recurring smooth scalp voltage topographies remaining stable for tens of milliseconds before rapidly reconfiguring (Lehmann et al., 1987; Michel and Koenig, 2018). A limited repertoire of such topographies can be identified with remarkable consistency across recordings and studies, providing a compact description of fast large-scale EEG dynamics and motivating their use to investigate physiological and pathological brain states and cognitive processes (Koenig et al., 2024; Michel and Bréchet, 2026).

### 1.1. Resting state EEG microstates and open methodological challenges

Microstate analysis can be broadly separated into two operations: finding representative topographies and using them to describe continuous EEG. Template estimation is commonly performed on maps sampled at peaks of global field power (GFP), where the electric field is strong and adjacent topographies are, on average, more stable (Zanesco, 2020). Candidate maps are then grouped using polarity-invariant clustering into a small set of representative templates (Pascual-Marqui et al., 1995; Michel and Koenig, 2018). The resulting templates are notably reproducible within and between datasets: different clustering procedures can yield highly consistent maps (Khanna et al., 2014; Kleinert et al., 2024), and cross-study aggregation reveals comparable templates across the resting-state literature (Koenig et al., 2024). Microstate templates are characteristically spatially smooth, which reinforced the adoption of spatial filtering to suppress isolated local fluctuations in clustered samples in dedicated software tools (Michel and Brunet, 2019).

Once templates have been estimated, they are used to construct a microstate sequence. In standard backfitting, each EEG map is compared with the available templates using polarity-invariant spatial similarity and assigned to its best match. Sequence construction is nevertheless not unique: labels may be assigned to every sample or propagated from GFP peaks, and postprocessing may subsequently reject, smooth or reassign samples according to correlation or duration criteria (Pascual-Marqui et al., 1995; Haydock et al., 2025). These choices matter because a weak assignment can have different causes. The scalp map itself may be poorly compatible with the broad geometry represented by microstates; alternatively, it may be coherent but insufficiently represented by the available template dictionary. A third case arises when two templates provide similarly strong matches, making winner-take-all assignment intrinsically ambiguous. Treating these situations identically can alter segment boundaries and therefore duration, occurrence, coverage and transition statistics. Although these measures can show good test-retest reliability under a fixed pipeline, the consequences of changing backfitting and postprocessing choices remain less systematically characterized (Michel and Koenig, 2018; Kleinert et al., 2024; Haydock et al., 2025).

Despite these methodological dependencies, microstate measures have shown reproducible associations with clinically and cognitively relevant states, including disorders of consciousness, the Alzheimer’s disease continuum, schizophrenia and attentional processes (Lassi et al., 2023; Toplutaş et al., 2024; De Pieri et al., 2025; Das et al., 2026; Ngo et al., 2026; Zanesco et al., 2026). However, the functional interpretation of individual microstate classes remains unsettled, and current perspectives emphasize converging evidence across methods and experimental contexts rather than assigning fixed functions to individual classes (Michel et al., 2024; Michel and Bréchet, 2026).

### 1.2. Continuous organization underlying resting-state microstates

The methodological challenges of postprocessing lead to a more fundamental question: in which space should microstate topographies and their transitions be represented to understand how assignment succeeds, becomes ambiguous or fails? Even more importantly, such a space should imply parallel evidence that are not contaminated by the choices used to generate the labels themselves. Recent work has approached this question by complementing discrete microstate sequences with continuous descriptions of the underlying EEG. Mishra et al. (2020) showed that expression of canonical microstates is not uniformly winner-take-all and that transitions can be gradual, with the degree of discreteness varying across the GFP range. From a different perspective, von Wegner et al. (2021) related recurrent microstate motifs to continuously rotating spatial phase patterns of resting-state alpha activity. Accordingly, recent methodological work has argued that microstate sequences should be cross-referenced against the continuous EEG dynamics from which they are derived (Haydock et al., 2025). Such approaches complement the well-established discrete microstate analysis by characterizing the underlying continuous dynamics.

A useful continuous representation should therefore satisfy several constraints. It should be defined independently of a particular clustering and backfitting solution, preserve the reproducible topographic organization on which microstate research is built, remain interpretable in terms of scalp-field shape, and explicitly retain the structure discrete labels do not capture. The literature itself provides a strong empirical lead. Koenig et al. (2024) organized template maps collected from 40 resting-state studies according to their pairwise topographic similarity, revealing reproducible regions and meta-microstate families in a low-dimensional data-driven space, as shown in Fig.1.

**Figure 1.**
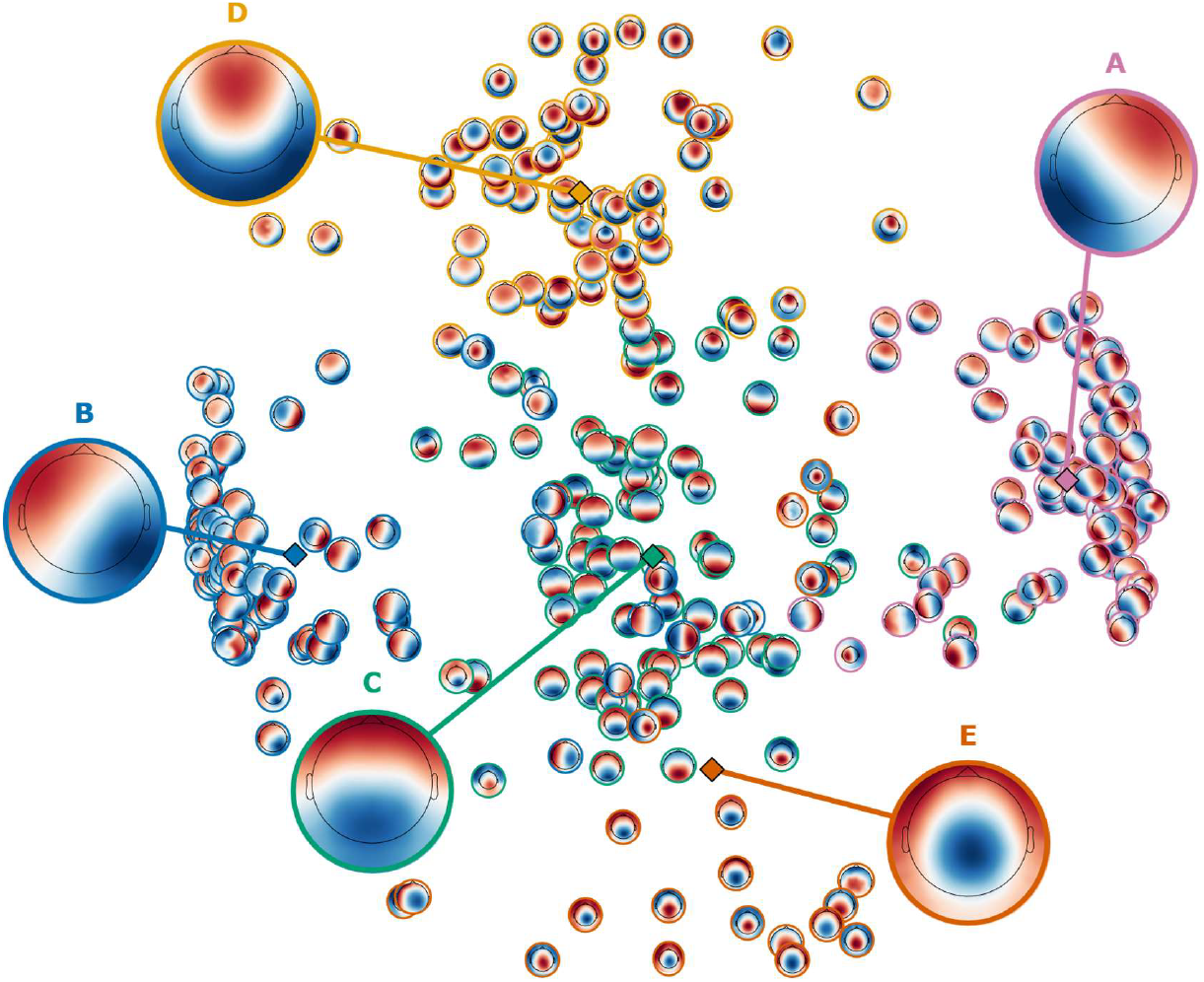
Literature templates and their gradient-like shapes. Group-wise microstate templates from the literature shown in the data-driven multidimensional scaling space obtained from the meta-clustering implemented in the MATLAB app MSTemplateExplorer (Koenig et al., 2024). Circle colors indicate the five-template meta-microstate solution derived in the app, with the center of each meta-cluster marked by a diamond.

As already apparent in early descriptions of microstate topographies, recurring configurations are dominated by broad smooth voltage gradients that the first publications also tried to describe via the lateralization of the vector connecting negative and positive centroids (Lehmann et al., 1987), although without distinguishing between linear and radial gradient patterns (Fig.1). We therefore asked whether this shared geometric property could provide a continuous, clustering-independent interpretable space in which canonical templates retain the same organization, and we hypothesized that the templates could be almost entirely described as linear or radial gradients with arbitrary lateralization.

### 1.3. The First-Order spherical haRMonic (FORM) hypothesis

On the spherical surface approximating the electrode montage, a simple three-dimensional family describing linear and radial gradients across arbitrary orientations is the First-Order spherical-haRMonic (FORM) subspace. The FORM can equivalently be represented by an electric dipole located at the center of the sphere: its three components along the transverse -right positive-, sagittal -frontal positive- and axial directions are linearly related to the voltage measured at each electrode and can therefore be estimated by least squares, together with an explicit residual. The dipole direction describes the orientation of the broad scalp gradient, while its magnitude, ρ, quantifies how strongly the original topography conforms to FORM space. This equivalent dipole is neither indicative of a single neural source nor a substitute for source reconstruction; it is simply a geometric representation of FORM topographies spanning linear and radial gradients with arbitrary lateralization. FORM therefore deliberately models a restricted broad-field family derived from characteristic properties repeatedly observed in microstate topographies. To validate this representation and test the consequences predicted by its formulation, we tested:

i. whether FORM selectively captures first-order spatial structure in controlled simulations and whether literature-derived, group-level and individual microstate templates strongly conform to it;
ii. whether FORM orientation provides an interpretable continuous coordinate system preserving the pairwise organization of known microstate maps;
iii. whether temporal topographic change can be decomposed into FORM and residual components, and whether change within FORM is predominantly driven by gradient orientation; and
iv. whether ρ provides a clustering-independent constraint on microstate assignment strength, defined as the correlation with the best-matching template.

The final prediction follows directly from the geometry of the representation. If microstate templates are strongly FORM-like, an EEG sample can only match them strongly if it is itself strongly FORM-like: in the ideal case, FORM conformity is therefore a necessary condition for strong microstate assignment. More strongly, under the ideal continuous FORM model, ρ is exactly the highest absolute correlation that a normalized EEG map can attain with any FORM topography. A finite template dictionary samples only a subset of FORM orientations and should therefore remain below this ceiling according to its directional coverage, while off-FORM structure in empirical EEG samples and templates produces departures from the ideal relation independently from the number and organization of template solutions. We consequently predicted that ρ would constrain winning-template correlation more directly than GFP and that increasing dictionary coverage would progressively reduce the average gap from the FORM ceiling. Together, these endpoints test whether FORM can provide a continuous, clustering-independent geometric reference for microstate topographies, their temporal evolution and the credibility of their discrete assignment, complementing the conventional microstate framework.

## 2. Results

### 2.1. FORM selectively captures first-order fields and strongly represents known microstate templates

#### 2.1.1. Simulated data with controlled spatial features

FORM selectivity was first tested in simulated voltage topographies whose spatial spectrum was fixed a priori, independently of any EEG data. We generated 1,000 maps under four conditions: pure first-order structure; spectra in which total degree-1 energy was five times degree-2 energy; spectra in which degree-1 energy was twice degree-2 energy; and spatial white noise, defined by equal expected energy per orthonormal spatial mode. Because the number of available modes increases with harmonic degree, the white-noise condition places progressively more total energy in higher spatial degrees. FORM reconstruction quality was quantified as the FORM modelling vector squared norm ρ^2^, the fraction of normalized map energy lying in the first-order subspace (Fig. 2).

**Figure 2.**
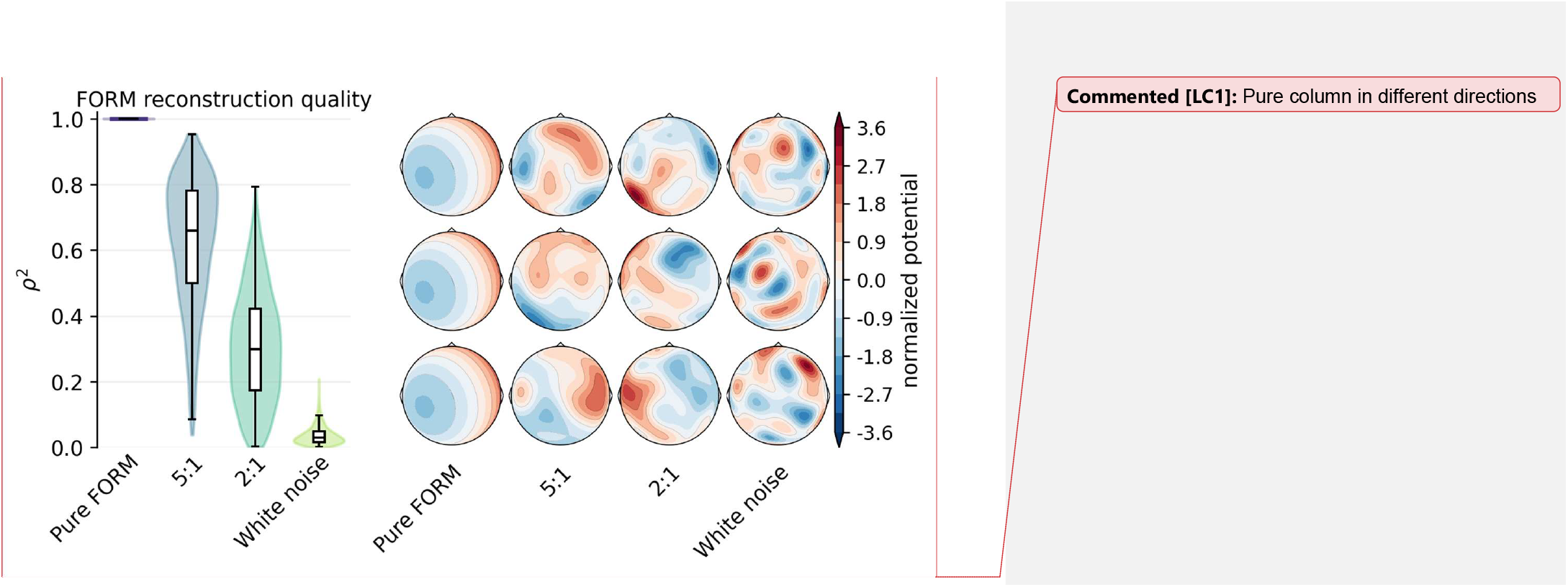
Controlled spatial-spectrum selectivity of FORM. The four spatial spectra were defined a priori as pure FORM; total degree-1 energy five times degree-2 energy; total degree-1 energy twice degree-2 energy; and spatial white noise, defined by equal expected energy per orthonormal spatial mode, so that higher harmonic degrees contribute more total energy since they contain more modes. (A) Distribution of FORM reconstruction quality *ρ*^2^ across 1,000 simulated maps per spectrum. (B) Three representative scalp topographies from the four spatial spectra on a 64-channel montage with normalized color scale.

As required by the orthogonal projection, pure first-order maps yielded ρ^2^ = 1. Introducing progressively flatter higher-order spectra reduced FORM conformity, with the 5:1 condition retaining more first-order structure than the 2:1 condition and white noise providing the lowest-conformity reference. The simulation therefore tests selectivity directly: FORM does not optimize a flexible model for each map but reports how much of that map belongs to the predefined first-order family.

#### 2.1.2. FORM representation of resting-state EEG meta-microstates and literature templates

Having established selectivity, we next asked whether the topographies conventionally identified as resting-state microstates occupy the same first-order family. For the five-template meta-microstate solution of Koenig et al. (2024), FORM reconstruction was near-complete, with *ρ*^2^ values of 0.992, 0.990, 0.998, 0.991 and 0.995 for A-E, respectively. We then projected the individual literature templates contributing to each meta-cluster, using the A-E correspondence adopted by Koenig et al. and aligned with the Custo et al. (2017) convention. Class-wise distributions are reported together with the corresponding meta-template values in Tab. 1, allowing the first-order structure to be assessed both at the level of the published maps and after meta-averaging.

**Table 1.** FORM-conformity of meta-templates and literature templates, considering meta-clustering with k=5 clusters. A full table with statistics computed in all cases from k=4, …, 8 and their visualization are available in Supplementary Materials (S1, S2).

| Map | Literature mean $\rho^2 \pm \text{SD}$ (N) | Literature median $\rho^2$ [q05,q95] (N) | Exact meta-template $\rho^2$ |
| --- | --- | --- | --- |
| A | 0.954 ± 0.029 (N=70) | 0.957 [0.948, 0.965] (N=70) | 0.992 |
| B | 0.946 ± 0.037 (N=78) | 0.958 [0.945, 0.962] (N=78) | 0.990 |
| C | 0.962 ± 0.026 (N=80) | 0.970 [0.963, 0.973] (N=80) | 0.998 |
| D | 0.960 ± 0.030 (N=61) | 0.966 [0.959, 0.971] (N=61) | 0.991 |
| E | 0.956 ± 0.030 (N=24) | 0.964 [0.954, 0.968] (N=24) | 0.995 |

#### 2.1.3. Microstate solution for the LEMON dataset and its FORM representation

The conventional clustering analysis provided the empirical solution against which FORM was tested, using the same preprocessed eyes-closed (EC) and eyes-open (EO) recordings and seven-criterion metacriterion as Zanesco (2020). For each recording, solutions with K = 1-12 were evaluated and the median of the seven criterion-specific optima defined the recording-level selected K. Centroids from the selected recording-level solutions were then pooled across EC and EO and reclustered for K = 1-15; application of the same metacriterion at group level selected K = 5, which was retained as the primary condition-insensitive LEMON dictionary. The pooled K = 5 solution was matched to the Koenig et al. (2024) A-E meta-templates by polarity-invariant Hungarian assignment, providing the canonical label reference for subsequent within-dataset correspondence. To characterize external correspondence, the resulting LEMON maps were also compared with the five Cartool 3.7 maps reported for the same dataset by Zanesco (2020). Fig. 3 reports the selection diagnostics and the three map sets, while complete similarity matrices are reported in Supplementary Figure S3. Values of GEV are reported in Supplementary Materials (S4).

**Figure 3.**
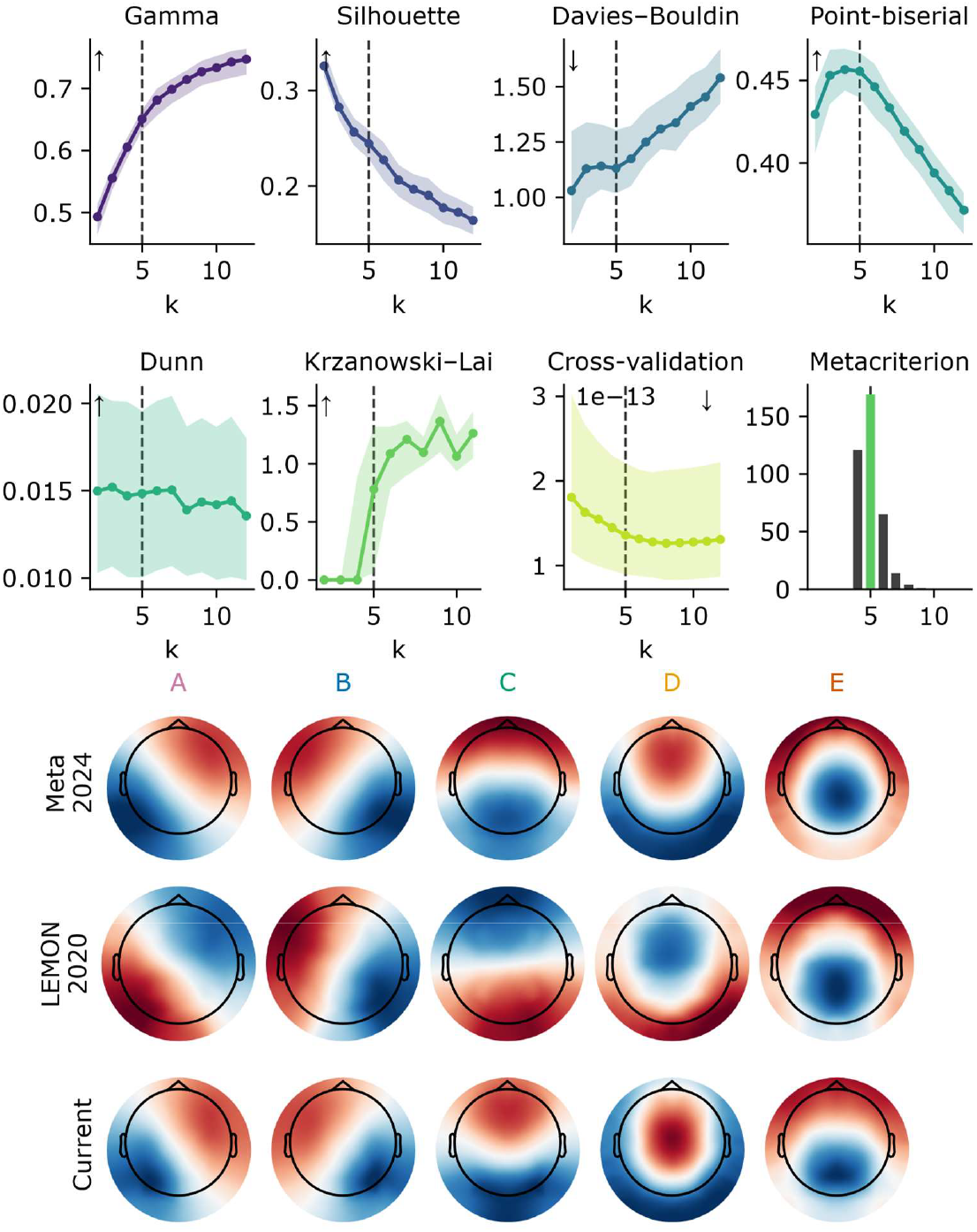
Conventional LEMON microstate solution and correspondence with external template sets. At the top, the seven clustering criteria evaluated independently across EC and EO recordings, shown as median and IQR, together with the distribution of recording-level K values selected by the seven-criterion metacriterion. Selected recording-level centroids were subsequently pooled and reclustered, with the same metacriterion selecting K = 5 at group level; the dashed vertical line marks this primary group solution. At the bottom, comparison of A-E topographies from the Koenig et al. (2024) meta-templates, the Cartool 3.7 LEMON solution reported by Zanesco (2020), and the present pooled LEMON K = 5 solution. The pooled LEMON solution was matched to the Koenig meta-templates by polarity-invariant Hungarian assignment and used as the canonical within-dataset reference.

FORM conformity of the LEMON templates was high at every aggregation level. For the pooled K = 5 group maps, ρ^2^ was 0.968, 0.966, 0.986, 0.985 and 0.972 for A-E, respectively. Condition-specific group maps were similarly first-order (ρ^2^ = 0.938-0.991), whereas subject-condition templates showed greater dispersion but retained median ρ^2^ values between 0.918 and 0.959 across label and condition (Fig. 4). Across the independently estimated K = 1-12 group dictionaries used in the ceiling analysis, every candidate template had ρ >= 0.978. The first-order representation therefore remained strong from literature meta-maps to empirical group templates and individual recording-level templates, while preserving measurable subject-level variability.

**Figure 4.**
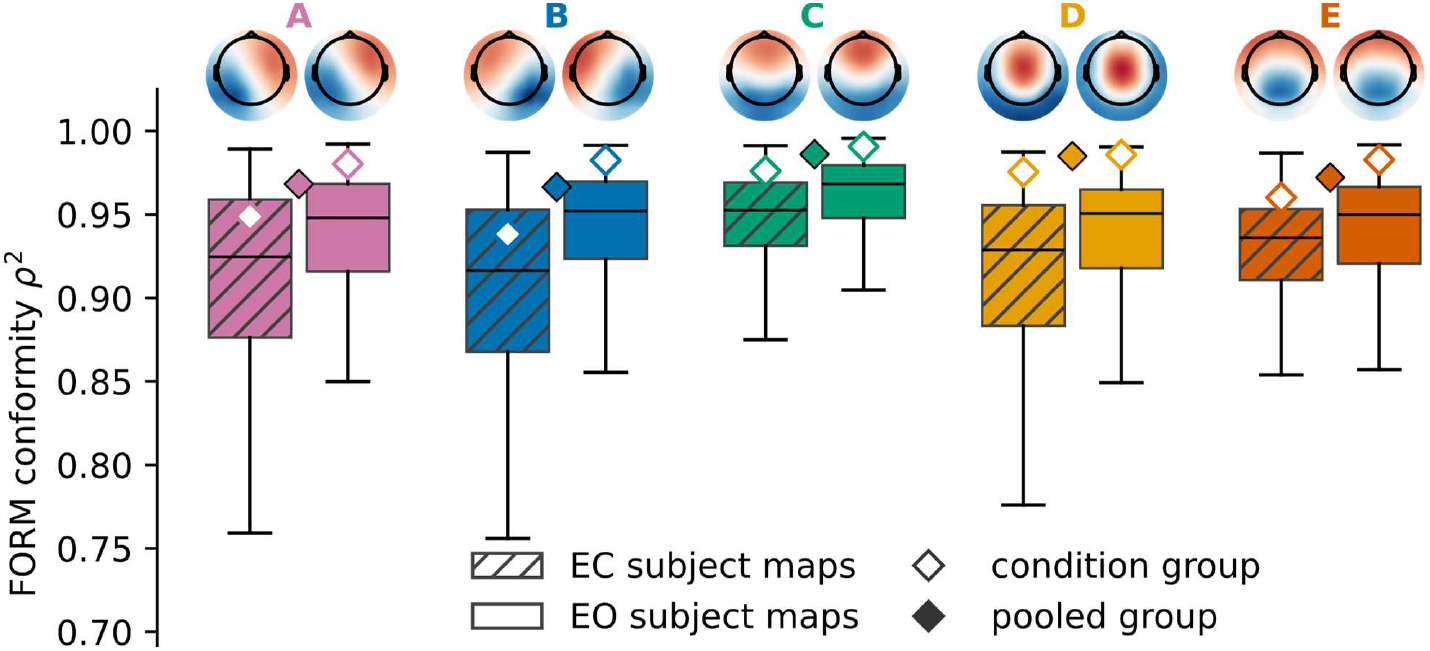
FORM conformity across LEMON template levels. Boxplots show ρ^2^ for subject-condition templates from 187 participants, separately for EC and EO within each microstate A-E. The condition-wise template maps, both matched to the reordered group-wise condition-insensitive templates, are shown at the top to show their similarity. Empty diamonds indicate FORM conformity of the condition-specific group templates shown at the top and filled diamonds indicate the conformity of the condition-insensitive K = 5 group templates.

### 2.2. FORM organizes microstates in an interpretable continuous space

Mapping literature-derived and empirical templates into polarity-canonical FORM orientation coordinates showed that the representation preserves an immediately readable relation between position in the space and visible map shape. In this coordinate system, azimuth captures lateralization, whereas elevation captures the continuum from near-linear anteroposterior gradients to more radial frontal or occipital field configurations. Templates lying near the equatorial band therefore correspond to predominantly linear gradients, while positive and negative elevations reflect increasing frontal and occipital radiality, respectively. This makes FORM coordinates directly interpretable in topographic terms, without relying on discrete class labels.

This organization was examined using the full set of literature maps included in the meta-microstate analysis of Koenig et al. (2024). Pairwise polarity-invariant angular distances between their FORM orientations closely tracked the pairwise topographic dissimilarities underlying the literature MDS (Spearman ρ = 0.987). This view-independent correspondence indicates that FORM preserves the relative organization of the literature templates without fitting the MDS embedding itself. The representation also provides an intuitive language to describe templates continuously, for example as a right-lateralized linear gradient microstate or as an occipital-radial gradient microstate, instead of referring only to canonical letter labels.

Within this organization, the lateralized canonical topographies corresponding to A and B occupied more localized regions of the space, whereas C, D and E were arranged more broadly along a low-lateralization anteroposterior continuum (Fig.5). This observation is reported here only as descriptive geometry. Its possible implications for the interpretation of canonical classes, for the continuous descriptions of microstates, and for methodological issues in clustering and assignment are presented in the Discussion.

**Figure 5.**
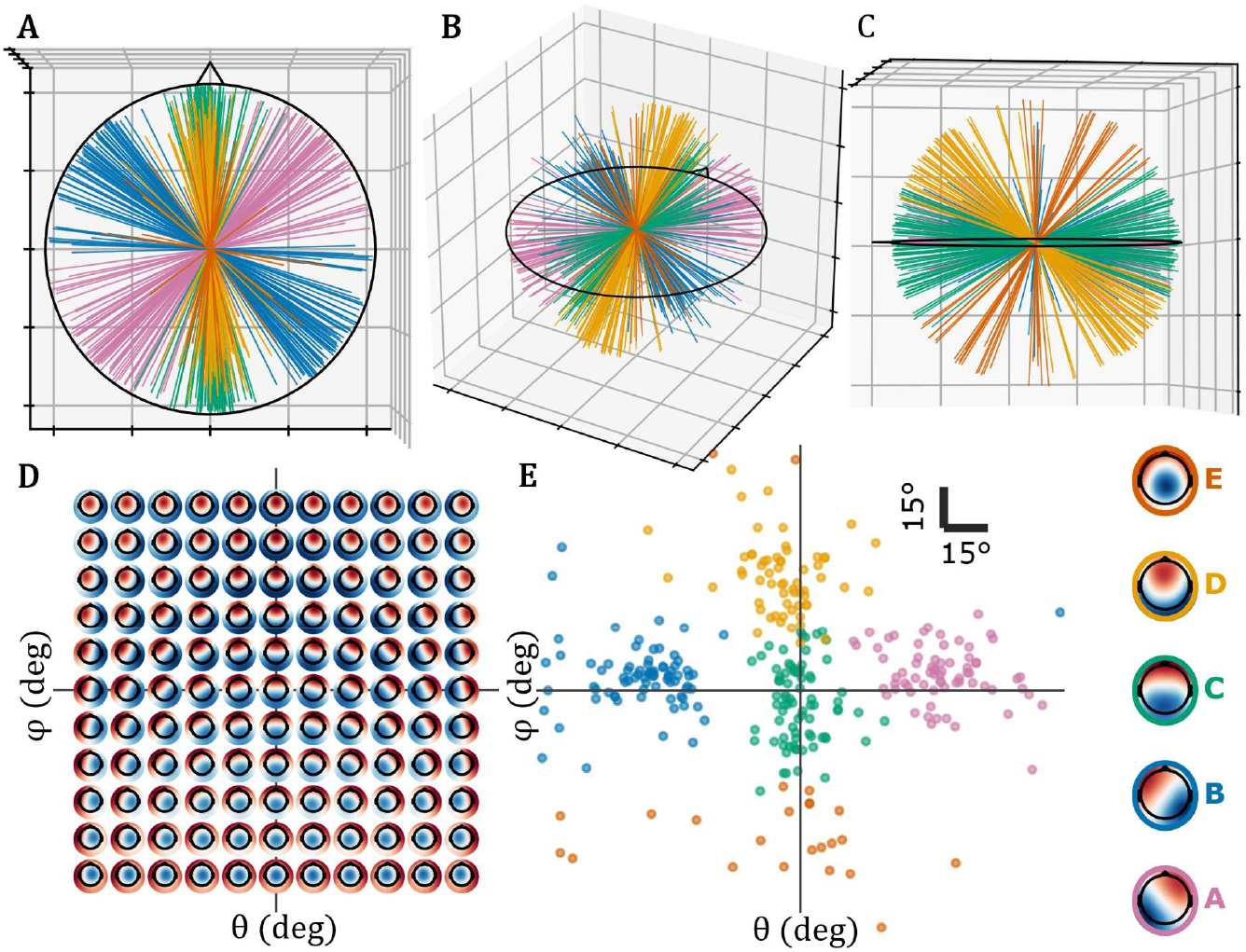
FORM orientation provides an interpretable continuous space for resting-state microstate topographies. At the top, coronal (A), isometric (B) and left sagittal (C) views of the FORM sphere for the canonical five-microstate solution, showing the corresponding Cartesian vectors in the three-dimensional FORM space. (D): grid of representative FORM topographies illustrating how visible scalp-map shape varies across the orientation space, from near-linear gradients around the equatorial band to increasingly frontal or occipital radial configurations at higher absolute elevations. (E): literature and meta-microstate templates projected into the two-angle polarity-canonical FORM space; map thumbnails, meta-cluster identities, and shape labels at the margins.

### 2.3. Temporal topographic change has separable FORM and off-FORM components

Topographic dissimilarity (TD), defined as the root mean square of the sample-to-sample change in scalp voltage topographies after normalization by instantaneous GFP, is classically known to decrease as GFP increases and provides one of the main rationales for sampling candidate maps at GFP peaks during subject-level clustering (Zanesco, 2020). We therefore asked whether the FORM representation could clarify why normalized topographies change over time. A map may change because its dominant FORM component changes orientation, consistent with previous observations of continuous topographic reorganization (von Wegner et al., 2021), or because part of the change lies outside the FORM subspace. We refer to these contributions as FORM dissimilarity and off-FORM dissimilarity, respectively.

From the FORM formulation, the squared TD between adjacent normalized maps can be decomposed exactly, up to negligible numerical closure error, into a FORM component and an off-FORM component. In turn, the squared FORM component can itself be decomposed into a term depending only on the change in FORM conformity, that is, the change in the modulus of the two FORM vectors, and a term depending on the change in FORM orientation, that is, the angular displacement between the two FORM vectors. We denote these two contributions as *TD*_*ρ*_ and *TD*_*ψ*_, respectively. This decomposition makes it possible to separate changes in how strongly a map conforms to the broad-field FORM family from changes in the orientation of that broad-field component.

Figure 6 summarizes these relationships for an exemplary EEG recording. We first reproduce the inverse TD-GFP power-law relationship previously described by Zanesco (2020), and then extend the same analysis to *TD, TD*_off-FORM_, *TD*_*ρ*_, and *TD*_*ψ*_. For simplicity, regressions are shown on TD rather than on squared TD. Because the analysis is performed in log-log space, using TD instead of *TD*^2^ only rescales the vertical axis and the regression coefficients by a factor of 2, without altering the qualitative structure of the fitted power laws. Alongside these examples, Table 2 reports group level summary metrics for each TD variant, the power law coefficient of the GFP relationship, and the coefficient of determination of the regression.

**Table 2.** Subject-level descriptive statistics and GFP power-law for decomposed topographic dissimilarity. For each measure, we report separately for EC and EO conditions the subject-level measures mean ± SD, the 5th and 95th percentiles, the mean ± SD of the scaled power-law regression coefficient relating the measure to GFP, the corresponding 5th and 95th percentiles, and the R^2^ of the relationship.

| Metric, condition | TD | $TD_{\text{FORM}}$ | $TD_{\text{off-FORM}}$ | $TD_{\psi}$ | $TD_{\rho}$ |
| --- | --- | --- | --- | --- | --- |
| TD, EC | $0.319 \pm 0.053$<br>[0.255, 0.427] | $0.233 \pm 0.031$<br>[0.193, 0.295] | $0.203 \pm 0.048$<br>[0.148, 0.291] | $0.220 \pm 0.029$<br>[0.183, 0.275] | $0.053 \pm 0.010$<br>[0.040, 0.071] |
| TD, EO | $0.358 \pm 0.066$<br>[0.276, 0.481] | $0.250 \pm 0.034$<br>[0.204, 0.310] | $0.239 \pm 0.063$<br>[0.164, 0.376] | $0.235 \pm 0.032$<br>[0.192, 0.290] | $0.059 \pm 0.012$<br>[0.044, 0.082] |
| Power-law $\beta$, EC | $-0.816 \pm 0.105$<br>[-0.969, -0.634];<br>$R^2 = 0.438$ | $-0.864 \pm 0.119$<br>[-1.046, -0.658];<br>$R^2 = 0.258$ | $-0.785 \pm 0.096$<br>[-0.900, -0.605];<br>$R^2 = 0.500$ | $-0.797 \pm 0.114$<br>[-0.977, -0.595];<br>$R^2 = 0.194$ | $-1.455 \pm 0.226$<br>[-1.784, -1.085];<br>$R^2 = 0.197$ |
| Power-law $\beta$, EO | $-0.833 \pm 0.093$<br>[-0.962, -0.678];<br>$R^2 = 0.445$ | $-0.889 \pm 0.098$<br>[-1.030, -0.711];<br>$R^2 = 0.246$ | $-0.806 \pm 0.094$<br>[-0.922, -0.660];<br>$R^2 = 0.497$ | $-0.814 \pm 0.090$<br>[-0.946, -0.651];<br>$R^2 = 0.178$ | $-1.540 \pm 0.241$<br>[-1.906, -1.114];<br>$R^2 = 0.191$ |

**Figure 6.**
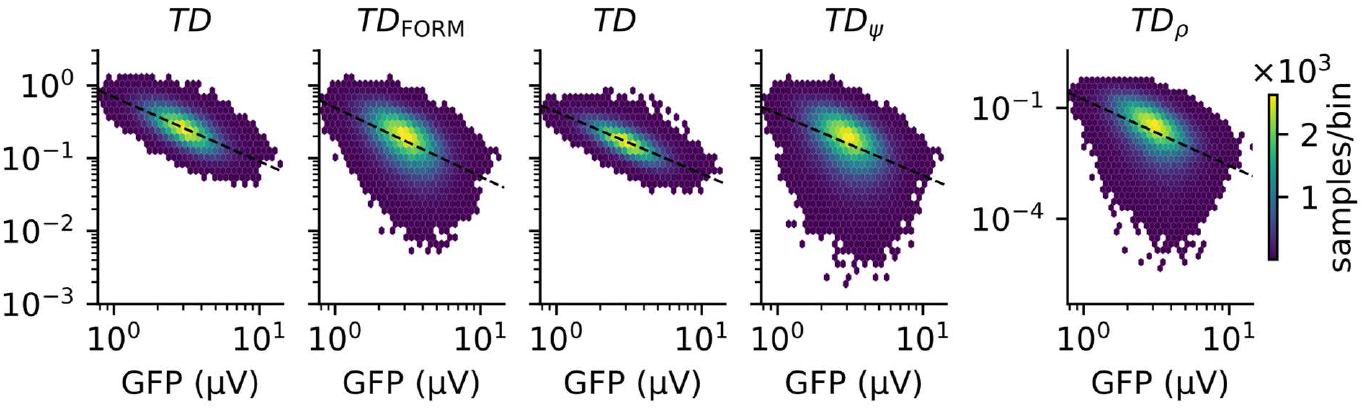
Temporal topographic change can be decomposed into FORM and off-FORM components. From left to right, panels show the log-log relationship between GFP and total topographic dissimilarity (TD); its decomposition into FORM and off-FORM components, *TD* and *TD*_off_; and the further decomposition of *TD* into changes in FORM conformity and orientation *TD*_*α*_, and *TD*_*ρ*_, respectively. *TD* closely follows *TD*_*α*_, as *TD*_*ρ*_ is approximately one order of magnitude smaller; *TD*_*ρ*_is therefore shown with a separate y-axis scale in the rightmost panel. Lines show the fitted log-log regressions. Density counts are expressed as thousands of samples per bin within the exemplary recording of the LEMON dataset, here subject 010214 in the EC condition.

Together, these results show that temporal topographic variation can be partitioned into interpretable geometric components. Within the FORM contribution, changes in orientation were markedly larger than changes in conformity: mean *TD*_*ψ*_ was approximately four times *TD*_*ρ*_ in both EC (0.220 vs 0.053) and EO (0.235 vs 0.059). Despite its smaller magnitude, the conformity component showed a steeper inverse GFP relationship, with power-law coefficients of *β* = −1.455 in EC and −1.540 in EO, compared with −0.797 and −0.814 for the orientation component. This supports the original microstate intuition of relatively stable broad field configurations that reorganize over time, while also showing that a substantial part of topographic change remains outside the FORM scaffold.

### 2.4. FORM conformity is a necessary condition of assignment strength

For a normalized EEG topography, *ρ* is exactly the maximum absolute correlation attainable with any unit topography in the continuous FORM family. Therefore, when candidate microstate templates are themselves strongly FORM-like, a sample can only be strongly assigned if it also has high FORM conformity. In this sense, FORM conformity is a necessary condition of assignment strength. This differs fundamentally from GFP: GFP quantifies field strength and is associated with topographic stability but places no analogous geometric constraint on how closely the normalized map can match a microstate template. Consistently, for the primary *K* = 5 solution, within-subject linear models based on GFP alone explained a median 21.7% of the variance in winning-template correlation, compared with 74.4% for *ρ* alone. Adding GFP to *ρ* produced only a small increase. The same relationship generalized across participants: pooled leave-one-subject-out predictions yielded *R*^2^ = 0.114 for GFP, *R*^2^ = 0.756 for *ρ*, and *R*^2^ = 0.760 for their combination. Full linear prediction results are reported in Supplementary Materials (S5).

For an ideal finite dictionary of FORM templates, this ceiling can be written as *r*_max_ = *ρ*cos(*δ*_*K*_), where *δ*_*K*_ is the angular distance to the closest template direction. Dividing assignment strength by *ρ* therefore isolates the directional-coverage term, motivating the normalized model *r*_max_/*ρ* = *β*_1_ + *ε*, in which *β*_1_ summarizes the average fraction of the continuous FORM ceiling reached by a finite dictionary. The empirical group templates closely satisfied the assumption underlying this interpretation: across the independently estimated *K* = 1 − 12 solutions, candidate-template *ρ* ranged from 0.978 to 0.995. Figure 7 illustrates the resulting ceiling relationship in the empirical sample space, while retaining the small departures expected because neither EEG samples nor empirical templates are perfectly restricted to FORM.

**Figure 7.**
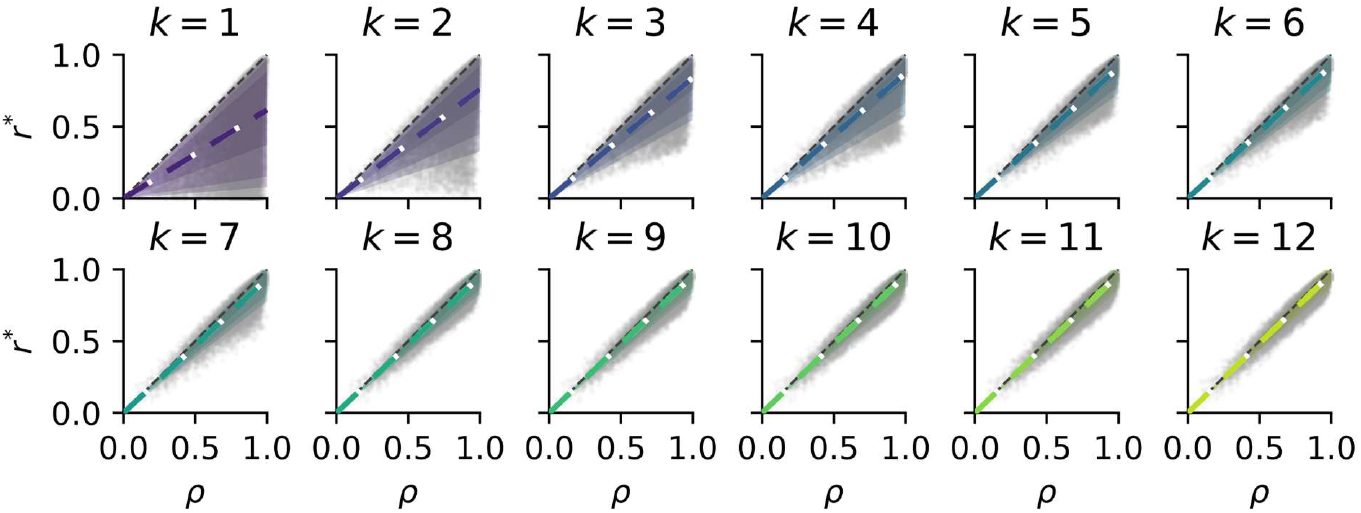
Relationship between FORM conformity and finite-dictionary assignment strength. For each independently estimated condition-insensitive group dictionary from *K* = 1to *K* = 12, winning-template correlation *r*_*K*_ is plotted against FORM conformity *ρ*. The grey dashed identity line, *r*_*K*_ = *ρ*, represents the continuous FORM ceiling. The colored dashed line shows the relation 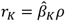 implied by the normalized ceiling-fraction model *r*_*K*_ /*ρ* = *β*_*K*_ + *ϵ*_*K*_. Here, 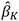 summarizes the average fraction of the continuous FORM ceiling reached by dictionary *K*. As *K* increases, the fitted relation progressively approaches the continuous ceiling, consistent with denser directional coverage of FORM space.

The normalized ceiling fraction increased systematically as more template directions were available (Fig. 8). Median subject-level *β*_*K*_increased from 0.613 at *K* = 1, to 0.911 at *K* = 5, and 0.970 at *K* = 12. The increase was monotonic across adjacent dictionary sizes for all 187 participants. At the same time, median raw-scale residual MAD decreased from 0.189 at *K* = 1, to 0.043 at *K* = 5, and 0.026 at *K* = 12. Thus, increasing dictionary size progressively reduced the average directional-coverage gap and brought finite-template assignment closer to the continuous FORM ceiling. Because each solution was estimated independently rather than constructed as a nested dictionary, this result describes an ordered cohort-level convergence and does not imply monotonic improvement for every individual EEG sample.

**Figure 8.**
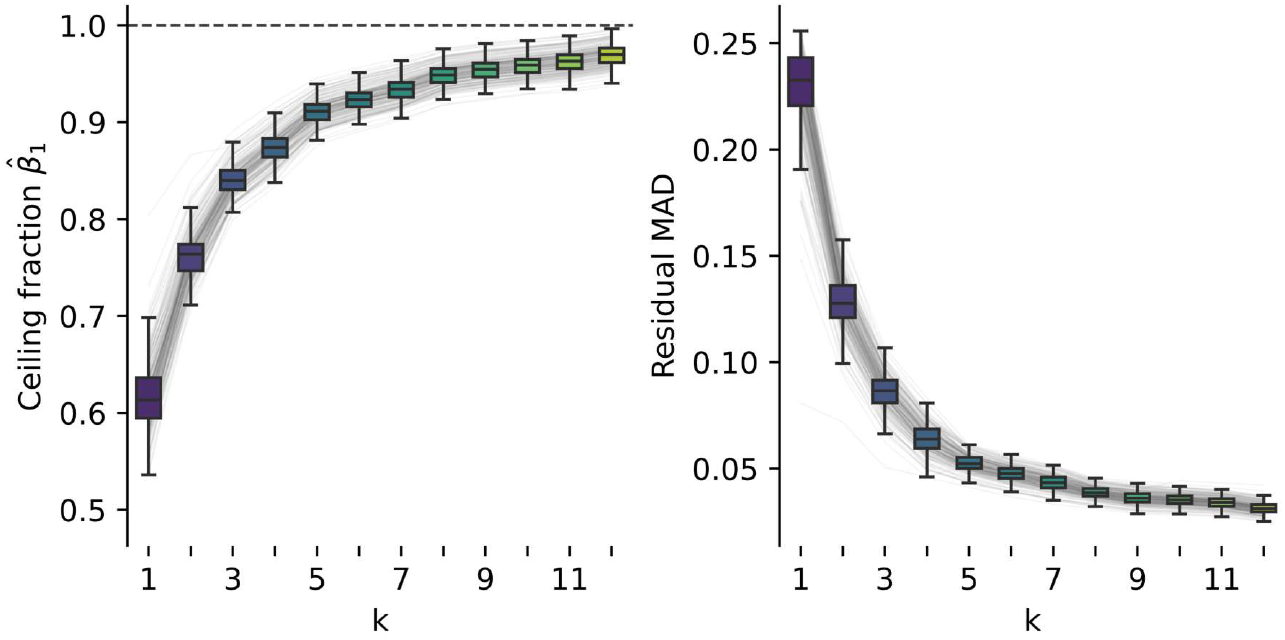
Finite microstate dictionaries progressively approach the continuous FORM ceiling. Left: subject-level estimates of the normalized ceiling fraction 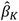 across independently estimated group dictionaries from *K* = 1to *K* = 12. Boxplots summarize the across-subject distribution, while grey trajectories connect estimates from the same participant; the horizontal dashed line marks the ideal continuous FORM ceiling, *β*_*K*_ = 1. Right: corresponding subject-level median absolute deviation (MAD) of the raw correlation-scale residual 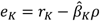. Together, the two panels show that increasing *K*improves average directional coverage while reducing the remaining deviation from the fitted ceiling relationship, without requiring adjacent solutions to be nested.

## 3. Discussion

FORM places resting-state microstate analysis in a reference geometry defined before clustering or backfitting. Broad, smooth and recurrent scalp configurations have been central to the microstate concept from its earliest formulation, while cross-study work has since shown a striking reproducibility of their canonical organization (Lehmann et al., 1987; Michel and Koenig, 2018; Koenig et al., 2024). With sensor geometry and reference convention fixed, the same first-order basis applies across cohorts and independently estimated microstate solutions. Empirical solutions can therefore be compared in a space that was not learned from those solutions. FORM and source reconstruction address almost opposite methodological aims, where source reconstruction expands sensor measurements towards models of neural sources, whereas FORM reduces the scalp field to a compact geometry intended for sensor-space comparison (Custo et al., 2017; Michel et al., 2024). The equivalent central dipole is indeed only an ideal equivalent of first-order field geometry, not a neural source model.

Field strength, field geometry and field change are separate quantities. GFP describes the strength of the instantaneous field; FORM conformity describes how closely its normalized shape belongs to the first-order family; topographic dissimilarity describes how that shape changes over time (Skrandies, 1990; Zanesco, 2020). We show that canonical resting-state templates are almost entirely contained in FORM, indicating that the first-order component closely captures the broad spatial structure principally targeted by conventional resting-state microstate analysis. The residual off-FORM structure is not synonymous with noise; it is spatial structure outside the family represented by those canonical maps. Separating the two components prevents higher-order variation from being folded into the same quantity used to describe the microstate-like field.

### 3.1. From weak correlation to assignment diagnosis

Conventional backfitting reduces several questions to the correlation with the winning template. Hard assignment has long been useful for constructing microstate sequences, but different procedures subsequently threshold, smooth, interpolate or reassign weak and short segments in different ways (Pascual-Marqui et al., 1995; Dinov and Leech, 2017; Haydock et al., 2025). FORM conformity adds information before any such decision: it defines the strongest match that the normalized map could attain anywhere in the continuous first-order family. Low conformity therefore identifies a map that cannot be closely represented merely by adding further FORM-like templates, whereas high conformity with only moderate winning correlation identifies an available first-order direction that the current dictionary represents poorly. In other terms, FORM conformity is a necessary condition of assignment strength. For strongly FORM-like templates, the fraction of the available FORM match reached by the winner becomes a measure of directional adequacy. Competition between two similarly good templates remains a third and separate uncertainty, which can still be described through winner-runner-up margins, probabilistic assignment approaches (Dinov and Leech, 2017), or via the elevation and azimuthal angle components of the distance to each of the best matching templates that are naturally induced by the FORM representation.

Increasing the number of microstate templates reduces the chance of leaving coherent FORM directions unrepresented, but directional adequacy is not itself a model-selection criterion. This distinction becomes relevant during sequence cleaning. A 25-ms interval with high FORM conformity but a poor template match may represent a coherent direction under-represented by the selected dictionary, whereas a 60-ms interval with low conformity reflects a departure from the first-order geometry itself. Duration rules cannot distinguish these cases (Haydock et al., 2025). A FORM-aware smoothing criterion could therefore penalize reassignment according to the GFP-weighted gap from the available FORM ceiling, 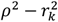, protecting highly conforming but under-represented intervals while allowing poorly conforming excursions to be smoothed more readily.

### 3.2. A continuous geometry underlying canonical microstates

The high FORM conformity of canonical templates supports the geometric observation from which the study started. This is not a low-dimensional representation learned from the microstate maps themselves. The FORM space is fixed by the first-order spherical harmonics, and the simulations show that its conformity decreases as more spatial structure lies in higher harmonic degrees. The strong conformity seen across literature, group and individual templates therefore means that the smooth gradients classically associated with resting-state microstates largely occupy this restricted spatial family.

The orientation coordinates then give this family a simple descriptive language. Azimuth mainly reflects lateralization, while elevation describes the progression from approximately linear gradients towards increasingly frontal or occipital radial configurations. Thus, two templates that would both be called microstate C, for example, can also be described in terms of a small angular displacement within the same broad field family. This remains a scalp-level description. The equivalent central dipole is only a representation of the three FORM coefficients and does not imply a single neural generator. The off-FORM residual instead represents spatial structure not captured by the first-order scaffold describing canonical resting-state microstate topographies. It should not be interpreted intrinsically as noise or irrelevant activity, since this residual may contain physiological, artifactual or other higher-order spatial structure.

### 3.3. A coordinate system for maps and trajectories

FORM direction gives every sufficiently conforming map an angular representation, therefore inducing an angular distance between maps automatically. Early microstate work already described broad topographic organization through the orientation connecting positive and negative field regions, while FORM extends that intuition to a continuous polarity-invariant geometry containing both linear and radial configurations (Lehmann et al., 1987). Azimuth and elevation therefore do more than provide compact coordinates: they state how far, and along which geometrically interpretable direction, one map has moved from another. Across the literature templates, pairwise FORM angular distances closely followed the pairwise topographic dissimilarities underlying the independently derived MDS organization. This does not establish a unique microstate space, but it shows that the predefined FORM geometry preserves the relative organization of literature topographies without being fitted to that organization.

Template comparison is an immediate application. TANOVA remains the inferential method for establishing whether multichannel topographies differ (Habermann et al., 2018; Nagabhushan Kalburgi et al., 2024). Once such a difference has been established, FORM can characterize it as a change in first-order conformity, an angular displacement within FORM, or both. A matched class can therefore be described quantitatively as becoming more radial or more lateralized relative to its reference, rather than only as differing topographically. Characterization of template differences is extremely relevant when comparing conditions or clinical groups because differences in the maps used for fitting can themselves propagate into apparent differences in temporal microstate parameters (Murphy et al., 2024). Angular changes remain scalp-level descriptions and do not necessarily identify altered neural sources, whose interpretation requires the separate evidence supplied by source imaging and multimodal studies (Britz et al., 2010; Custo et al., 2017; Milz et al., 2017).

Continuous trajectories extend the angular description from templates to the EEG between labels. Previous work supports both structured microstate dynamics and substantial continuity between canonical configurations (Van De Ville et al., 2010; Mishra et al., 2020; Creaser et al., 2021; von Wegner et al., 2021). These alternatives can be examined directly in FORM space: if microstates correspond to metastable regions, trajectories should accumulate around restricted orientations and cross rapidly between them; if the underlying evolution is smoother, trajectories should move continuously through broader regions whose discrete identities arise partly from the chosen dictionary. FORM provides coordinates in which occupancy, angular velocity and departures from the first-order family can distinguish these patterns while remaining linked to scalp topography (Haydock et al., 2025).

### 3.4. Practical scope and limitations

Spatial filtering provides a further, narrower use. Spatial smoothing is already present in established topographic and microstate workflows, including CARTOOL, to reduce local spatial fluctuations before topographic analysis (Brunet et al., 2011; Michel and Brunet, 2019). FORM projection offers a linear alternative whose retained spatial family is explicitly defined and, according to the present results, closely matches canonical resting-state microstate geometry. It should not be regarded as a generally superior EEG filter. FORM is better viewed as a principled spatial restriction when the target is specifically sensor-space microstate analysis; wherever ordinary spatial filtering would be inappropriate, including analyses that require the discarded spatial information, FORM projection offers no special reason to apply it.

Clustering independence does not imply montage independence. Electrode positions and coverage, reference, interpolation and sphere fitting determine the discrete first-order basis, while electrode density is already known to influence the reliability of conventional microstate estimates (Zhang et al., 2021). The conditioning analysis also showed that the FORM basis remains well behaved across progressively sparser representative montages; the 64-channel montage used here had *κ*(*M*) = 1.288, close to the isotropic limit of 1.

Empirical validation remains limited to healthy adult resting-state EC and EO recordings together with published resting-state templates. Clinical, developmental, task-related and artifact-rich data may have different conformity and residual distributions, so no universal conformity threshold follows from the present cohort (Kleinert et al., 2024; Michel and Bréchet, 2026). Low conformity only states that a scalp field is poorly represented by the broad first-order geometry characteristic of the resting-state templates examined here; the residual may contain physiological, pathological or artifactual structure that requires separate interpretation. The ideal directional account further assumes templates that themselves lie very close to FORM, while empirical residual components can locally perturb the expected relationship. Independently estimated solutions with different numbers of templates are not nested, so convergence towards the FORM ceiling is a cohort-level tendency rather than guaranteed improvement for every sample. Continuous EEG observations are also strongly autocorrelated, requiring participant-level inference or explicit treatment of temporal dependence rather than treating the sample cloud as independent observations.

### 3.5. Conclusion

FORM adds a question that comes before deciding which microstate wins: how much of the scalp field belongs to the geometry that canonical microstates actually occupy? Because canonical templates are almost entirely first-order, FORM conformity sets a geometric ceiling on how strongly such templates can match, while FORM orientation states where the field lies within that space. This separates off-FORM structure from incomplete dictionary coverage, turns template differences into interpretable angular displacements, and opens the discrete sequence back into a continuous trajectory. FORM does not replace discrete microstates; it makes the geometry beneath their labels explicit, measurable and continuous.

## 4. Materials and methods

### 4.1 Established microstate formalism

We provide here a brief overview of conventional resting state EEG microstate analysis focused on aspects useful for the understanding of the FORM space approach. The reader is referred to established methodological and review literature for further details (Pascual-Marqui et al., 1995; Michel and Koenig, 2018; Nagabhushan Kalburgi et al., 2024; Haydock et al., 2025).

#### 4.1.1 Scalp voltage, global field power and topographic dissimilarity

Let ***x***(*t*) ∈ *R*^*N*^ denote the scalp-voltage vector for an EEG with *N* channels. In the following we always assume average-referencing, i.e. ***x***(*t*) is zero-mean. The spatial standard deviation takes the name of Global Field Power (GFP) and is thus defined as:

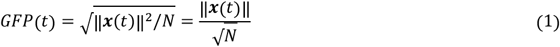

In microstate analysis, since label assignment is based on correlation, the object of interest is actually the spatially z-scored potential, i.e. ***x***(*t*)/*GFP*(*t*). In the case of the FORM approach, we use the L2-normalized potential ***z***(*t*) to keep equations more readable:

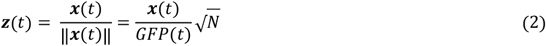

In the temporal dimension, the main quantity is the Topographical Dissimilarity (TD), defined as the sample-to-sample root-mean-square backward change in EEG topography, normalized to be independent of instantaneous field strength (Skrandies 1990, Zanesco 2020):

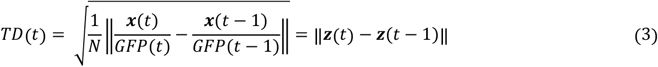

The TD is thus naturally expressed as the norm of the backward finite difference in time of the L2-normalized EEG topography. The widespread choice of using GFP peaks of subject-level clustering comes from an empirical relationship linking TD and GFP by a scaled power law, usually verified by linear regression in logarithmic space (Zanesco 2020):

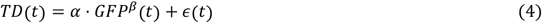

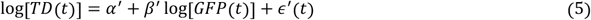

The *β* coefficient is known to be consistently negative even at subject level (Zanesco 2020), providing solid evidence that samples with strong field are more topographically stable in time.

#### 4.1.2 Microstate templates, assignment and postprocessing

Conventional microstate templates are estimated by polarity-invariant clustering of representative scalp maps, commonly sampled at local GFP maxima. Let ***b***(*k*) ∈ *R*^*N*^ denote a centered unit-norm microstate template from a dictionary containing *K* maps. The polarity-invariant match of sample ***z***(*t*) to template k is the correlation:

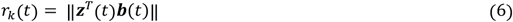

and backfitting procedure assigns the EEG sample of interest to a winning template with maximum correlation *r*^*\**^(*t*) = *max*_*k*_[*r*_*k*_(*t*)] the template label *k*^*\**^(*t*) = *argmax*_*k*_ [*r*_*k*_(*t*)]. The backfitting procedure does not by itself consider the correlation margin, which can instead be independently reported as the difference between *r*^*\**^(*t*) and the second largest correlation. The main quality control metric for backfitting is the Global Explained Variance (GEV), which can be defined at group-level, subject-level, and for the backfitting procedure and indicate the variance explained by the templates, considering the variance respectively of the individual maps clustered at group-level, of the candidate EEG samples used in subject-level clustering, and of all backfitted EEG samples in a recording. The first two quantify adequacy of the clustering procedure, while backfitting GEV is usually considered a feature of microstate analysis.

While labeling of microstate templates is essentially arbitrary, comparability across literature requires association of similar templates to a same common name: given the impressive reproducibility of the last two decades of resting state topographies in the literature, similarity is usually established by a Hungarian matching algorithm as implemented in the MICROSTATELAB toolbox using either the solutions found in Custo et al. (2017) or Koenig et al. (2024) following the lexicographic order of Custo et al. (2017) such that A is commonly a linear right-lateralized anteroposterior gradient, B a linear left-lateralized anteroposterior gradient and so on.

Instantaneous backfitting produces a sequence of winning microstate labels across EEG samples. Depending on the analysis pipeline, the resulting sequence may subsequently be modified by rejecting samples below a minimum template correlation, favouring temporally coherent labels through local smoothing, or reassigning segments shorter than a predefined minimum duration (Pascual-Marqui et al., 1995; Haydock et al., 2025). A commonly used form of the Pascual-Marqui smoothing iteratively assigns each sample by minimizing a loss penalizing topographic misfit and favouring temporal continuity. Some pipelines instead fit only GFP peaks and propagate the resulting labels to the intervening samples. These operations affect the temporal boundaries of the final sequence and therefore the derived duration, occurrence, coverage and transition measures. In the analyses below, instantaneous topographic matching is kept separate from these subsequent sequence-level operations.

### 4.2 FORM representation

FORM represents each centered, L2-normalized EEG topography by its projection onto a three-dimensional first-order subspace defined by the sensor geometry, the fitted electrode sphere and the chosen EEG reference. This subspace corresponds to the family of centered first-order, dipole-like scalp fields. The FORM representation is therefore defined a priori, independently of the EEG data and of any microstate clustering solution.

#### 4.2.1 FORM as a least squares problem

The sensor geometry is conventionally expressed in cartesian coordinates centered on the best head-fitting sphere. The axis convention depends on the software ecosystem: in EEGLAB (MATLAB) (Delorme and Makeig, 2004) and MNE-Python (Gramfort et al., 2013), for example, the first coordinate of an electrode *x*_*i*_ represents the sagittal and transverse coordinate, respectively, while the center is the same and the last coordinate *z*_*i*_ is in both cases the axial coordinate. Regardless of the specific convention, let ***R*** ∈ *R*^*N*×3^ denote the montage matrix whose i-th row is the cartesian positions of the *N* electrodes relative to the sphere center.

For an electric dipole with moment ***p*** located at the sphere center, the potential at electrode position *r*_*i*_ in a homogeneous isotropic volume conductor with is proportional to:

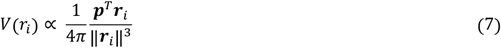

via the inverse of the conductivity *σ*. Writing the same equation over all electrodes gives the linear model for ***z***(*t*) and consequently the least squares dipole estimate:

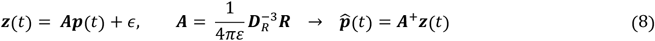

where ***A***^+^ denotes the pseudoinverse of ***A*** and we have defined the diagonal radius matrix ***D***_*R*_:

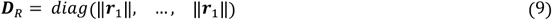

The fitted scalp field 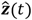 is then ***AA***^+^***z***(*t*). In this direct physical formulation, the design matrix contains both electrode direction and electrode-specific radial weighting through 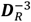. In the ideal case in which all electrodes lie at the same radius *r*, ***D***_*R*_ = ***rI***_*N*_, so *A* differs from *R* only by a global scalar. The three Cartesian coordinates then correspond exactly to the three real spherical harmonics of degree one, defining the three-dimensional FORM subspace.

#### 4.2.2 FORM as an orthonormal transform

The least-squares formulation above provides a physically direct three-dimensional design matrix *A*, but its coefficients retain both the radial weighting and the metric induced by the electrode geometry. FORM can instead be formulated directly on the spherical surface and expressed as an orthogonal transform of the scalp topography.

The first step is to remove the physical radial scale from the montage, which is irrelevant since we are treating normalized topography and not physical voltage. We define the normalized montage matrix ***U*** ∈ *R*^*N*×3^ as:

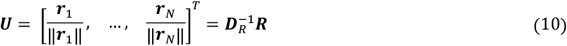

When all electrode positions lie on the same spherical surface, ***D***_*R*_ = ***rI***_*N*_, so *U* and the dipole design matrix ***A*** of the previous formulation differ only by a global scalar and therefore span the same three-dimensional field family. For real montages, the two formulations are not strictly identical: the normalization defining *U*removes electrode-to-electrode differences in radius and retains only the angular sensor geometry that defines the spherical FORM space.

Each column of ***U*** represents the scalp topography produced by one of the three components of the dipole. For an arbitrary coefficient vector ***a***(*t*) the corresponding first-order field is therefore ***v***(*t*) = ***Ua***(*t*). However, the measured EEG topography ***z***(*t*) is expressed after average referencing. Thus, the model prediction must be expressed under the same reference before comparing it with ***z***(*t*). Common average referencing can be expressed as the matrix ***H***_*CAR*_ such that the re-referenced field ***v***_*CAR*_(*t*) = ***H***_*CAR*_***v***(*t*) is zero-mean:

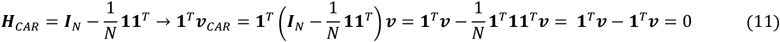

The columns of ***H***_*CAR*_***U*** therefore describe the same three first-order fields after average-referencing, so that any linear combination of an arbitrary dipole is zero mean as the normalized topography ***z***(*t*).

The three columns, however, are not generally orthogonal. Even after radial normalization, a finite EEG montage does not sample the sphere isotropically, so its three Cartesian directions can have different norms and non-zero pairwise inner products. We collect this residual montage geometry in the 3 × 3 Gram matrix

**Commented [LC2]:** Posso dire che ortononormalizzo e basta

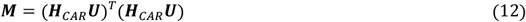

is therefore the Gram matrix of the FORM design before orthonormalization and plays the same role for its conditioning as ***A***^T^***A*** in the direct least-squares formulation. We remove this metric by symmetrically normalizing the three directions using ***M***^−1/2^ and defining the orthonormal basis ***Q***:

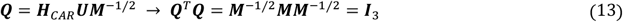

Finally, ***Q*** is the orthonormal basis of the FORM family spanned by ***H***_*CAR*_***U***. After finding ***Q***, the FORM representation of an arbitrary topography ***z***(*t*), the reconstructed FORM topography and the error or off-FORM topography are:

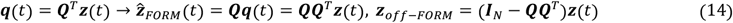

The stepwise construction makes explicit the three operations required to obtain the FORM transform from the original montage: radial normalization through 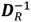, average referencing through *H*_*CAR*_, and orthonormalization of the resulting three-dimensional basis through ***M***^−1/2^. Each operation therefore has a distinct geometric or signal-processing role rather than being absorbed into a single fitted transformation as in the simpler least-squares derivation.

The orthogonal formulation also gives a direct interpretation of the resulting exact decomposition of Eq. 14 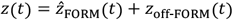 where the two components are orthogonal:

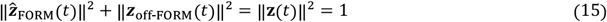

Moreover, because ***Q*** has orthonormal columns, the mapping from FORM coefficients to the FORM subspace is an isometry:

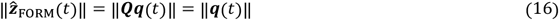

and projection of an arbitrary sensor-space topography onto FORM is non-expansive: the norm of the projection, which we term *ρ*(*t*) or FORM conformity, is thus lower than the norm of the normalized topography, which is unitary by definition. *ρ*(*t*) lies between 0 and 1 and its square is exactly the fraction of normalized topographic energy explained by the FORM projection:

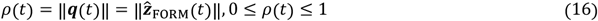

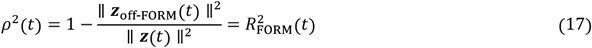

where the final equality follows because ***z***(*t*) is centred and unit-norm. Thus, *ρ*^2^ is simultaneously the squared length of the FORM projection and the coefficient of determination of its orthogonal reconstruction.

We therefore use the spherical orthogonal formulation throughout the following analyses. It coincides with the direct central-dipole least-squares formulation up to a global scale in the ideal common-radius case, while for real montages it deliberately removes radial variability before average referencing and orthonormalization.

The stability of the three-dimensional fit depends on electrode geometry. Differences in electrode radius alter the relative sensor weights, while directional anisotropy can leave one or more spatial directions weakly sampled and is typically the more important source of ill-conditioning in standard EEG montages. To quantify montage anisotropy, we used the spectral condition number of the pre-orthonormalization Gram matrix ***M*** = (***H***_CAR_ ***U***)^T^(***H***_CAR_ ***U***), *κ*(***M***) = *λ*_max_(***M***)/*λ*_min_ (***M***), with values close to 1 indicating approximately isotropic spatial sampling. As a practical reference, *κ*(***M***)was 1.182, 1.288, 1.351, and 1.432 for representative 128-, 64-, 32-, and 19-channel montages, respectively. The montage used in the present analyses corresponds to the 64-channel case (*κ* = 1.288).

#### 4.2.3 FORM conformity and orientation

For a centered unit-norm topography ***z***, FORM coordinates are ***q*** = ***Q***^T^***z***, and the FORM conformity *ρ* is the norm of the orthogonal FORM projection (Eq.16). Thus, the unit vector:

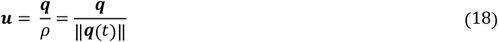

Represents FORM orientation independently of conformity; since resting state EEG microstates are polarity invariant, ***u*** and −***u*** represent the same FORM orientation. The same orientation can also be described in spherical coordinates, and polarity invariance is enforced selecting, for an arbitrary orientation, the vector lying in the anterior half of the spherical space approximating the head: in other terms, the vector is inverted when its sagittal component *u*_*y*_ < 0. We thus define the elevation angle *Φ*:

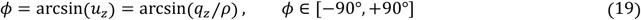

And the azimuthal angle *θ*:

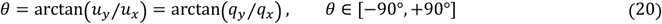

Moreover, when *u*_*y*_ = 0, *θ* takes the sign of *u*_*x*_. Because polarity canonicalization restricts the vector to the anterior half-sphere, no additional quadrant correction is required so that both angles are defined between −90 deg and +90 deg.

Elevation describes the progression from approximately linear to increasingly radial field configurations. Near-null values correspond to predominantly linear gradient-like topographies, whereas increasingly positive and negative elevations indicate progressively more frontal and occipital radial configurations, respectively. Azimuth describes lateralization: negative values correspond to left-lateralized configurations, values close to 0^°^ to approximately non-lateralized configurations, and positive values to right-lateralized configurations. For strongly FORM-conforming maps, (*θ, Φ*) therefore provides an almost complete description of the dominant smooth gradient-like scalp geometry.

#### 4.2.4 Decomposition of topographic dissimilarity

For adjacent centered unit-norm scalp maps ***z***_*t*−1_ and ***z***_*t*_, we defined their sample-to-sample topographic dissimilarity TD in Eq.3. Using the orthogonal decomposition 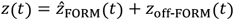, squared topographic change decomposes exactly as

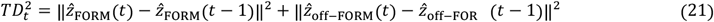

We denote the two square-root components as *TD*_FORM_ and *TD*_off-FORM_, respectively. The FORM component can be decomposed further using the norm and orientation defined in Section 4.2.3, where 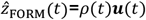. Thus, the norm of the difference in Eq.21 can be written as:

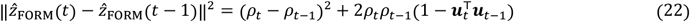

We therefore defined:

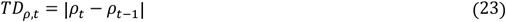

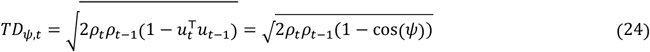

*TD*_*ρ*,t_ quantifies changes in FORM conformity and *TD*_*ψ,t*_ quantifies changes in FORM orientation, where *ψ* is the angle between two consecutive orientations.

#### 4.2.5 FORM conformity as a geometric ceiling for assignment strength

Let *F* = col(**Q**) denote the three-dimensional FORM subspace and let **z** be a centered unit-norm topography. Its orthogonal FORM projection is 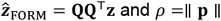. For any unit-norm topography **m** ∈ *F*,

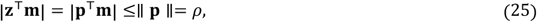

with equality attained by 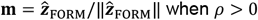. Therefore,

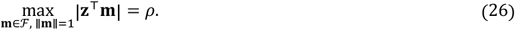

FORM conformity is consequently the exact continuous-family ceiling on polarity-invariant spatial correlation with a unit topography contained in FORM.

For a finite ideal dictionary with unit FORM directions **u**_*k*_, let *δ*_*K*_ be the smallest polarity-invariant angular distance between the sample orientation **u** and the available template directions. The best attainable correlation is then

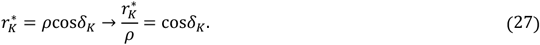

The normalized ratio therefore isolates finite directional coverage from sample conformity. As the dictionary samples FORM orientation more densely, the average value of *r*^*\**^/*ρ* is expected to increase towards one. However, empirical templates are not perfectly confined to FORM. Writing a unit template as

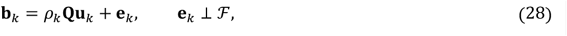

and the sample as **z** = *ρ***Qu** + **e**, their signed sensor-space similarity is

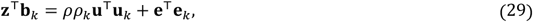

with

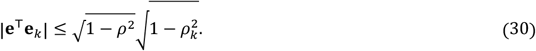

Thus, the ideal multiplicative relation is approached when candidate templates are strongly FORM-conforming, while off-FORM structure introduces an additional perturbation that cannot be removed simply by increasing directional coverage. Thus, we chose a normalized model:

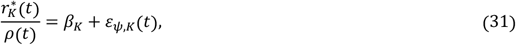

where *β*_*K*_ is the participant-level mean fraction of the continuous FORM ceiling reached by dictionary *K* and *ε*_*ψ,K*_ represents sample-to-sample variation in directional coverage.

### 4.3 Validation using literature templates and controlled spatial simulations

Before applying FORM to a real dataset, we evaluated the representation using only externally defined microstate topographies and synthetic fields with controlled spatial structure. These analyses tested whether established resting-state microstate topographies occupy the predefined FORM family, whether their pairwise topographic organization is preserved by FORM orientation, and whether FORM conformity selectively decreases when spatial energy is moved towards higher-order structure.

#### 4.3.1 Literature-derived microstate maps and external geometric organization

We first evaluated FORM using the resting-state microstate material compiled in the MSTemplateExplorer MATLAB app by Koenig et al. (2024). This included the meta-template solutions containing four to eight meta-clusters and the individual literature templates contributing to each meta-cluster. We focused on the five-cluster solution to match the five-state solution previously reported for the LEMON dataset by Zanesco (2020). Class labels followed the correspondence adopted by Koenig et al. and the lexicographic convention of Custo et al. (2017). We additionally considered the five Cartool 3.7 templates reported by Zanesco (2020) as a historical solution obtained from the same dataset subsequently analyzed empirically.

Each map was represented in its documented sensor geometry, centered, L2-normalized and projected onto FORM. For every template we extracted FORM conformity *ρ*, its explained-energy equivalent *ρ*^2^, the three-dimensional FORM coordinates **q**, and the polarity-canonical orientation coordinates (*θ, Φ*). Conformity quantified how completely established microstate topographies were represented within the predefined first-order subspace, while the angular coordinates were used to visualize their topographic organization.

We then tested whether the data-driven organization of the literature templates was preserved in FORM geometry. For the 313 templates, we compared the original pairwise topographic dissimilarities used to construct the MDS with the polarity-invariant angular distances between the corresponding unit FORM orientations. For orientations ***u***_*i*_ and ***u***_*j*_, FORM distance was defined as 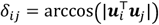. Correspondence between the two distance matrices was summarized descriptively by Spearman rank correlation across their upper triangles, containing each unique template pair once. Because this comparison uses pairwise distances rather than MDS coordinates, it is independent of the arbitrary orientation and viewing angle of the three-dimensional MDS embedding.

#### 4.3.2 Controlled spatial-spectrum simulations

FORM selectivity was tested independently of empirical EEG and of the literature templates using synthetic topographies with predefined spatial spectra. Synthetic fields were generated in a montage-adapted orthonormal spatial-harmonic basis with degrees *ℓ* = 1,…,8. Let *Q*_*ℓ*_ contain the *n*_*ℓ*_ retained orthonormal modes of degree *l*. Within each condition, every retained mode at degree *ℓ* was assigned the same expected energy proportional to *w*_*ℓ*_ = *ℓ* ^−*α*^, so that the expected total energy of degree *ℓ* was *p*_*ℓ*_ ∝ *n*_*ℓ*_ *ℓ* ^−*α*^.

Four spectra were used: pure FORM, for which only degree 1 was retained; *α* = log_2_(25/3) ≈ 3.06, giving *p*_1_/*p*_2_ = 5; *α* = log_2_(10/3) ≈ 1.74, giving *p*_1_/*p*_2_ = 2; and *α* = 0, corresponding to equal expected energy per retained orthonormal mode. Thus, in the white-noise condition, total degree energy was proportional only to the number of retained modes at that degree. Each resulting topography was subsequently L2-normalized before FORM projection.

### 4.4 Validation using open resting-state EEG data

#### 4.4.1 LEMON resting-state EEG dataset

We analyzed the openly available Leipzig Study for Mind-Body-Emotion Interactions (LEMON) resting-state EEG dataset (Babayan et al., 2019), following the organization used in previous large-sample microstate analyses of the same cohort (Zanesco, 2020). Resting EEG was acquired in 16 alternating one-minute eyes-closed (EC) and eyes-open (EO) blocks beginning with EC. We used the preprocessed EC and EO recordings available for this analysis and retained only participants with valid paired conditions at the expected 250 Hz sampling rate. The final cohort comprised 187 participants and 374 recordings.

The analysis sensor space was harmonized to a fixed 63-channel montage consisting of the 61 LEMON scalp channels plus Fpz and FCz as in Zanesco (2020). The exact target channel set was: Fp1, Fpz, Fp2, AF7, AF3, AF4, AF8, F7, F5, F3, F1, Fz, F2, F4, F6, F8, FT7, FC5, FC3, FC1, FCz, FC2, FC4, FC6, FT8, T7, C5, C3, C1, Cz, C2, C4, C6, T8, TP7, CP5, CP3, CP1, CPz, CP2, CP4, CP6, TP8, P7, P5, P3, P1, Pz, P2, P4, P6, P8, PO7, PO3, POz, PO4, PO8, PO9, O1, Oz, O2, PO10, Iz. Target channels absent from an individual recording were added and interpolated from the available scalp sensors, channels outside the target set were removed, channels were reordered identically across recordings, and the data were re-referenced to the common average. All subsequent sensor-space calculations used this fixed channel order and reference.

#### 4.4.2 Standard microstate analysis

For each EC and EO recording separately, GFP was computed across the average-referenced montage and local GFP maxima were identified. Maps at GFP peaks were clustered with polarity-invariant modified k-means as implemented in Pycrostates 0.6.1 (Férat et al., 2022). Recording-level solutions were fitted independently for *K* = 1, …, 12 using 100 initializations, a maximum of 300 iterations and convergence tolerance 10.

Recording-level model selection used seven criteria: Gamma, Silhouette, Davies-Bouldin, point-biserial, Dunn, Krzanowski-Lai and cross-validation (Zanesco, 2020). For each recording, the optimum K was determined separately for each criterion, with criterion optima considered for K ≥ 4. The recording-level metacriterion was defined as the median of the seven criterion-specific optimal K values, and the centroids from the resulting recording-specific selected solution were contributed to group pooling. Selected recording-level centroids from both EC and EO recordings were pooled into a condition-insensitive group dataset and clustered again with independent modified k-means solutions for K = 1-15, using 200 initializations, a maximum of 300 iterations and a convergence tolerance of 10^-6. The same seven-criterion metacriterion was then applied to the group-level solutions and selected K = 5, which was retained as the primary pooled group dictionary. Independent group solutions for K = 1-12 were retained for the finite-dictionary coverage analysis; these solutions were estimated independently and were not nested extensions of one another.

The primary five pooled group templates were aligned to canonical A-E labels by polarity-invariant Hungarian matching against the five Koenig et al. (2024) meta-templates, following the Custo et al. (2017) convention. This labelled pooled solution was then retained as the reference for within-dataset correspondence. Recording-level K = 5 templates were matched to the pooled group templates for class-specific analyses. Condition-specific group solutions were obtained by pooling the K = 5 recording-level centroids separately for EC and EO, reclustering each pool at K = 5 using the same modified k-means procedure and matching the resulting templates to the pooled group reference. The Cartool 3.7 maps reported by Zanesco (2020) were retained as an external historical comparison.

#### 4.4.3 FORM metrics computation

For every sample, the average-referenced scalp map was centered and L2-normalized and projected with the fixed FORM basis of the analysis montage. We retained the three FORM coordinates **q**(*t*), conformity *ρ*(*t*) = ∥ **q**(*t*) ∥, explained-energy fraction *ρ*^2^(*t*), polarity-canonical static angles (*θ*(*t*), *Φ*(*t*)), the reconstructed FORM component and the complementary off-FORM component. The same transform was applied to literature, recording-level and group-level template maps represented in the corresponding sensor geometry.

For consecutive valid samples within the same continuous block, total topographic dissimilarity was decomposed according to Section 4.2.4 into *TD*_FORM_, *TD*_off-_, *TD*_*ρ*_ and *TD*_*ψ*_. Transitions crossing block boundaries or other identified temporal discontinuities were excluded. Analyses of temporal change additionally required valid non-zero normalized topographies at both endpoints. Numerical closure errors for the FORM/off-FORM and radial/angular decompositions were retained as quality-control quantities.

FORM conformity was summarized for the Koenig meta-templates, individual literature templates, the pooled LEMON group templates, condition-specific EC and EO group templates, recording-level templates aligned according to the hierarchy described above, and every candidate template in the independently estimated group dictionaries used for the ceiling analysis.

#### 4.4.4 Prediction of instantaneous assignment strength

To determine whether FORM conformity constrains instantaneous assignment strength more directly than field amplitude, the response was defined as the winning absolute correlation *r*^*\**^(*t*) with the condition-insensitive pooled group *K* = 5 dictionary. This target was computed before any smoothing, duration constraint or correlation-based rejection.

Within each participant, EC and EO samples were pooled and three ordinary least-squares models were fitted: GFP alone, *ρ* alone, and GFP plus *ρ*. GFP was expressed in microvolts, and predictors were standardized within the fitting data. The models were descriptive at this stage and were summarized by *R*^2^, RMSE and correlation between observed and predicted assignment strength. To keep the samplewise analysis computationally bounded and participant-balanced, at most 1,500 deterministically and evenly spaced samples were retained per participant.

Generalization across participants was evaluated by leave-one-subject-out prediction. At each fold, predictor means and standard deviations were estimated only from the training participants, the linear model was fitted on the training set, and predictions were generated for the held-out participant. Performance was summarized both for each held-out participant and from the concatenated held-out predictions using *R*^2^, RMSE and observed-predicted correlation. The comparison was intended to distinguish the geometric prerequisite represented by *ρ* from the more indirect association between GFP and topographic stability.

#### 4.4.5 Decomposed topographic dissimilarity relationship with GFP

The established GFP-TD relationship was evaluated separately for total TD and for each component of the FORM decomposition. For every participant and condition, ordinary least-squares regressions were fitted on finite positive-valued samples in log-log coordinates,

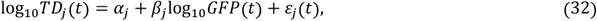

for *j* ∈ {total, FORM, off-FORM, *ρ, ψ*}. GFP was expressed in microvolts. Regressions were performed on the unsquared TD scale for direct comparability with conventional TD analyses; using squared TD would preserve the log-log relation while rescaling the slope and intercept.

For each component and separately for EC and EO, we report participant-level mean TD and fitted *β* coefficients summarized across participants by the mean, standard deviation, 5th and 95th percentiles, together with the coefficient of determination of the regression.

#### 4.4.6 Modeling FORM conformity as the upper bound for assignment strength

Finite-dictionary convergence was evaluated using the independently estimated condition-insensitive group dictionaries for *K* = 1, …, 12. For each participant and 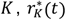 was computed as the maximum absolute sensor-space correlation between the normalized EEG sample and the available group templates.

The model was the normalized ceiling-fraction model introduced in Section 4.2.5, with the multiplicative coefficient 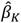 as the participant-level estimate:

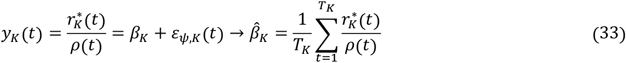

Under the ideal finite-FORM relation, 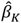 represents the mean fraction of the continuous FORM ceiling reached by dictionary *K*, obtained by averaging the samplewise ratio 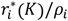. Under the ideal finite-FORM relation this corresponds to the average directional-coverage term cos*δ*_*K*_. Residual dispersion in the primary model was summarized from 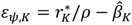.

## 6. Declarations and reproducibility statements

### CRediT authorship contribution statement

**LC:** Conceptualization, Methodology, Software, Formal analysis, Visualization, Writing - original draft, Writing - review and editing. **PL:** Methodology, Validation, Writing - original draft, Writing - review and editing. **CMO:** Supervision, Writing - review and editing. **AM:** Supervision, Funding acquisition, Writing - review and editing. All authors reviewed and approved the final version of the manuscript.

### Ethics and secondary-data statement

This study consisted exclusively of secondary analyses of previously collected data and involved no new recruitment, intervention, or data collection from human participants. Ethical approval and informed consent for the original data collection were obtained by the investigators responsible for the respective source datasets, as reported in the original studies and associated dataset documentation. No additional ethical approval was required for the present secondary analysis.

### Data availability

The data analysed in this study were obtained from the previously existing preprocessed dataset at ftp://ftp.gwdg.de/pub/misc/MPI-Leipzig_Mind-Brain-Body-LEMON/ (Zanesco, 2020). No new primary data were collected for this study.

### Code availability

The code used to perform the analyses and generate the results reported in this study is available at https://github.com/Leonardo-Corsi/microstates-form

## Acknowledgements

The authors have no acknowledgments to declare.

## Declaration of AI-assisted technologies in the writing process

During the preparation of this manuscript, the authors used generative AI tools for language proofreading and editing with the sole aim of improving grammatical correctness, syntactic clarity, and consistency of exposition. Generative AI was not used as a substitute for scientific reasoning, methodological decisions, data analysis, or interpretation of the results. All AI-assisted outputs were thoroughly reviewed in full by the authors, without exception, and the authors take full responsibility for the content of the manuscript.

