## Supplementary material for "Resting state EEG microstates: is a new FORM all you need?": S1

**Supplementary Table S1. FORM conformity of literature-derived microstate templates across meta-clustering solutions from K = 4 to K = 8.** For each meta-cluster, FORM reconstruction quality is reported as $\rho^2$ for the individual literature templates assigned to that cluster, summarized as mean ± SD and median [5th and 95th percentiles], together with the number of contributing templates (N). The final column reports the exact $\rho^2$ of the corresponding meta-template derived by Koenig et al. (2024). For every k, solutions are aligned to Custo et al. (2017), so the same map is consistent across different k values.

| K | map | Literature $\rho^2$ Mean ± SD (N) | Literature $\rho^2$ Median [$q_{05}$,$q_{95}$] (N) | meta-template $\rho^2$ (exact) |
|---|---|---|---|---|
| 4 | A | 0.953 ± 0.029 (N=71) | 0.956 [0.948, 0.965] (N=71) | 0.992 |
| | B | 0.945 ± 0.038 (N=80) | 0.958 [0.945, 0.962] (N=80) | 0.990 |
| | C | 0.962 ± 0.024 (N=94) | 0.967 [0.964, 0.972] (N=94) | 0.998 |
| | D | 0.961 ± 0.029 (N=68) | 0.966 [0.961, 0.972] (N=68) | 0.991 |
| 5 | A | 0.954 ± 0.029 (N=70) | 0.957 [0.948, 0.965] (N=70) | 0.992 |
| | B | 0.946 ± 0.037 (N=78) | 0.958 [0.945, 0.962] (N=78) | 0.990 |
| | C | 0.962 ± 0.026 (N=80) | 0.970 [0.963, 0.973] (N=80) | 0.998 |
| | D | 0.960 ± 0.030 (N=61) | 0.966 [0.959, 0.971] (N=61) | 0.991 |
| | E | 0.956 ± 0.030 (N=24) | 0.964 [0.954, 0.968] (N=24) | 0.995 |
| 6 | A | 0.953 ± 0.031 (N=63) | 0.956 [0.948, 0.965] (N=63) | 0.992 |
| | B | 0.945 ± 0.038 (N=63) | 0.958 [0.944, 0.963] (N=63) | 0.991 |
| | C | 0.962 ± 0.026 (N=79) | 0.969 [0.961, 0.973] (N=79) | 0.998 |
| | D | 0.961 ± 0.030 (N=62) | 0.966 [0.960, 0.973] (N=62) | 0.991 |
| | E | 0.956 ± 0.030 (N=24) | 0.964 [0.955, 0.968] (N=24) | 0.995 |
| | F | 0.951 ± 0.028 (N=22) | 0.956 [0.946, 0.964] (N=22) | 0.968 |
| 7 | A | 0.953 ± 0.031 (N=60) | 0.957 [0.948, 0.967] (N=60) | 0.992 |
| | B | 0.945 ± 0.038 (N=62) | 0.958 [0.943, 0.963] (N=62) | 0.991 |
| | C | 0.961 ± 0.027 (N=64) | 0.969 [0.958, 0.973] (N=64) | 0.997 |
| | D | 0.956 ± 0.034 (N=39) | 0.963 [0.955, 0.971] (N=39) | 0.990 |
| | E | 0.955 ± 0.031 (N=22) | 0.963 [0.952, 0.966] (N=22) | 0.995 |
| | F | 0.951 ± 0.028 (N=22) | 0.956 [0.946, 0.964] (N=22) | 0.968 |
| | G | 0.968 ± 0.018 (N=44) | 0.971 [0.962, 0.975] (N=44) | 0.995 |
| 8 | A | 0.953 ± 0.031 (N=60) | 0.957 [0.948, 0.967] (N=60) | 0.992 |
| | B | 0.945 ± 0.037 (N=65) | 0.957 [0.944, 0.963] (N=65) | 0.991 |
| | C | 0.961 ± 0.028 (N=63) | 0.969 [0.958, 0.973] (N=63) | 0.997 |
| | D | 0.956 ± 0.034 (N=39) | 0.963 [0.955, 0.971] (N=39) | 0.990 |
| | E | 0.961 ± 0.016 (N=19) | 0.965 [0.955, 0.966] (N=19) | 0.995 |
| | F | 0.953 ± 0.027 (N=15) | 0.960 [0.944, 0.967] (N=15) | 0.973 |
| | G | 0.968 ± 0.018 (N=44) | 0.971 [0.962, 0.975] (N=44) | 0.995 |
| | H | 0.933 ± 0.051 (N=8) | 0.946 [0.906, 0.972] (N=8) | 0.970 |
