## Supplementary material for "Resting state EEG microstates: is a new FORM all you need?": S2

**Supplementary Figure S2. FORM conformity of literature-derived templates across meta-clustering solutions from K = 4 to K = 8.** For each meta-cluster and each solution size, boxplots summarize the distribution of FORM conformity ($\rho^2$) across the individual literature templates assigned to that cluster. Jittered points show individual templates, and diamond markers indicate the exact $\rho^2$ of the corresponding meta-template. Labels A-H follow the canonical cluster ordering used in each K solution.

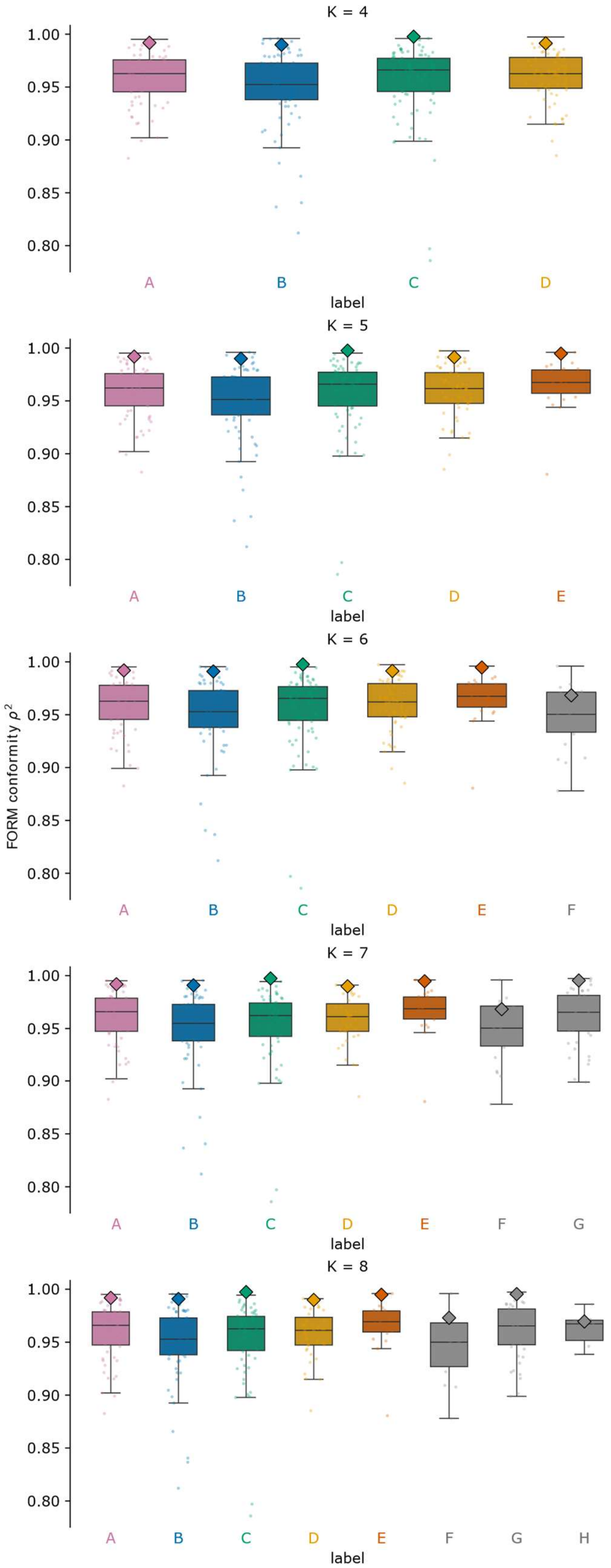
