## Supplementary material for "Resting state EEG microstates: is a new FORM all you need?": S3

**Supplementary Figure S3. Similarity of the current LEMON group solution to external five-template reference sets.** Absolute spatial correlations (|r|) are shown between the present K = 5 group templates and the Koenig et al. (2024) meta-microstates (top) and the Cartool 3.7 LEMON templates reported by Zanesco (2020) (bottom). Rows correspond to the current A-E templates and columns to the external reference templates; cell values report the pairwise absolute correlations used to assess cross-solution correspondence.

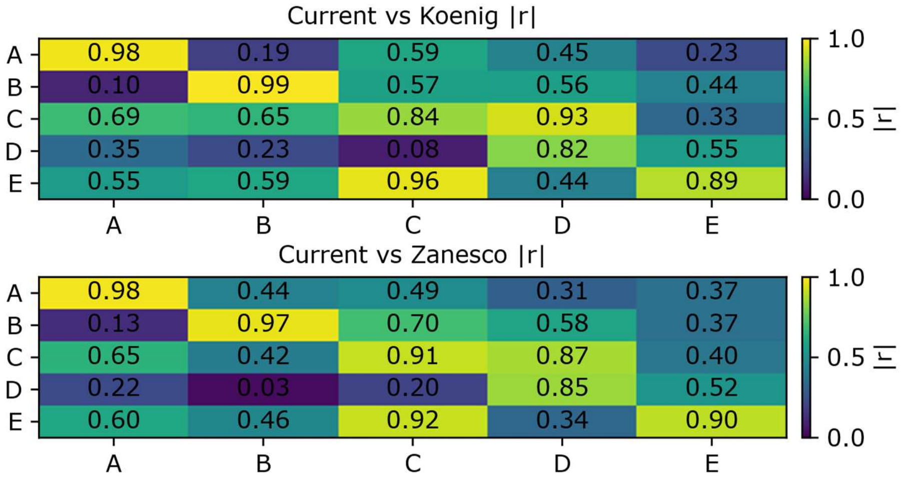
