## Supplementary material for "Resting state EEG microstates: is a new FORM all you need?": S4

**Supplementary Table S4. Global explained variance at the three microstate-analysis levels**. Subject-level clustering GEV is summarized across recordings from each recording's own GFP-peak candidate maps; group-level clustering GEV summarizes the fit of the K = 5 group solution to subject-level templates; backfitting GEV summarizes the fit of the same group solution to continuous EEG. A-E columns are microstate-specific contributions after the hierarchical label correspondence described in Methods, and sum to Total within each row. Values are reported separately for EC and EO, but GEV is computed at group-level and backfitting from the group-wise condition-insensitive templates.

| GEV level | Condition | A | B | C | D | E | Total |
|---|---|---|---|---|---|---|---|
| Subject-level clustering | EC | 0.116 | 0.127 | 0.232 | 0.113 | 0.162 | 0.751 |
| Subject-level clustering | EO | 0.114 | 0.128 | 0.196 | 0.103 | 0.158 | 0.700 |
| Group-level clustering | EC | 0.143 | 0.160 | 0.276 | 0.075 | 0.170 | 0.824 |
| Group-level clustering | EO | 0.159 | 0.161 | 0.215 | 0.087 | 0.186 | 0.807 |
| Backfitting | EC | 0.091 | 0.106 | 0.263 | 0.075 | 0.138 | 0.673 |
| Backfitting | EO | 0.100 | 0.112 | 0.182 | 0.081 | 0.154 | 0.630 |
