## Supplementary material for "Resting state EEG microstates: is a new FORM all you need?": S5

**Supplementary Table S5. Performance of assignment-strength linear prediction based on global field power and FORM conformity.** Linear models predicted the winning-template assignment correlation from GFP alone, FORM conformity (ρ) alone, or both predictors. Within-subject performance was evaluated on each recording's sampled observations. Leave-one-subject-out (LOSO) performance was evaluated by fitting each model on all other recordings and predicting the held-out recording. Values are mean ± SD across 187 recordings; r is the Pearson correlation between observed and predicted assignment strength, $R^2$ is the coefficient of determination, and RMSE is the root mean squared error. The GFP-only model showed limited generalization, whereas ρ accounted for most of the explainable variation; adding GFP yielded a small additional improvement.

| **Predictor model** | **Within-subject r** | **Within-subject $R^2$** | **Within-subject RMSE** | **LOSO r** | **LOSO $R^2$** | **LOSO RMSE** |
|---|---|---|---|---|---|---|
| *GFP* | *0.484 ± 0.057* | *0.238 ± 0.051* | *0.128 ± 0.011* | *0.484 ± 0.057* | *0.099 ± 0.128* | *0.140 ± 0.016* |
| *ρ* | *0.878 ± 0.022* | *0.772 ± 0.038* | *0.070 ± 0.004* | *0.878 ± 0.022* | *0.766 ± 0.038* | *0.071 ± 0.004* |
| *GFP + ρ* | *0.883 ± 0.021* | *0.780 ± 0.038* | *0.068 ± 0.004* | *0.882 ± 0.021* | *0.775 ± 0.039* | *0.069 ± 0.004* |
